# Advances in the recovery of haplotypes from the metagenome

**DOI:** 10.1101/067215

**Authors:** Samuel M. Nicholls, Wayne Aubrey, Kurt de Grave, Leander Schietgat, Christopher J. Creevey, Amanda Clare

## Abstract

High-throughput DNA sequencing has enabled us to look beyond consensus reference sequences to the variation observed in sequences within organisms; their haplotypes. Recovery, or assembly of haplotypes has proved computationally difficult and there exist many probabilistic heuristics that attempt to recover the original haplotypes for a single organism of known ploidy. However, existing approaches make simplifications or assumptions that are easily violated when investigating sequence variation within a metagenome.

We propose the metahaplome as the set of haplotypes for any particular genomic region of interest within a metagenomic data set and present Hansel and Gretel, a data structure and algorithm that together provide a proof of concept framework for the recovery of true haplotypes from a metagenomic data set. The algorithm performs incremental haplotype recovery, using smoothed Naive Bayes — a simple, efficient and effective method.

Hansel and Gretel pose several advantages over existing solutions: the framework is capable of recovering haplotypes from metagenomes, does not require a priori knowledge about the input data, makes no assumptions regarding the distribution of alleles at variant sites, is robust to error, and uses all available evidence from aligned reads, without altering or discarding observed variation. We evaluate our approach using synthetic metahaplomes constructed from sets of real genes and show that up to 99% of SNPs on a haplotype can be correctly recovered from short reads that originate from a metagenomic data set.

## Introduction

Genomic research has gone beyond finding a consensus sequence to represent a species in favour of understanding the true genetic diversity that exists within populations (Gibbs et al., 2003). High-throughput sequencing has allowed us to look further than consensus variation across populations of a species and reconstruct the genetic variants that characterizes each individual (haplotype).

Haplotype recovery approaches (also known as haplotype assembly) attempt to reconstruct haplotypes given a set of DNA fragments from a population (Lancia et al., 2001). Typically such approaches rely on the availability of reference sequences for the species under investigation. However, microbial communities consist of a large number of organisms for which no references exist and cannot at present be cultured *in vitro*. This has led to approaches that isolate and sequence DNA directly from the environment (metagenomics).

Microbial communities contain many organisms that co-exist in competition for the available resources and these communities represent an untapped wealth of genetic diversity. Information on this diversity is typically lost through current *de novo* analysis pipelines that make assumptions about ploidy and the source of the DNA. The majority of computational approaches assume reads originate from a single individual.

This impacts our ability to isolate the full repertoire of enzymes from microbial communities. If solved, this would advance exploitation of industrially relevant enzymes for the refinement of biofuels, production of plastics, scrub oil from water and even identify new classes of antibiotics (Zhang and Kim, 2010).

We would like to determine the collection of haplotypes for any genomic region of interest such as a particular enzyme, which we define as the **meta-haplome**.

### Assembly and pseudo-references

An assembly of reads from a metagenome can act as a pseudo-reference in the absence of a fully assembled genome. However, the pseudo-reference is a consensus of the read information and cannot represent the true haplotypes present. Furthermore, the pseudo-reference may not exist in nature and not constitute a viable enzyme. Once we have constructed the pseudo-reference, we discard the only evidence of the real haplotypes, the reads themselves.

Most assemblers are designed for single species genomes, are optimised to remove low level variation, and aim to produce a single sequence. Metagenome assemblers improve this with techniques for management of large data and correcting poorly assembled contiguous sequences (contigs) (Namiki et al., 2012). However they do not aim to solve the problem of recovering haplotypes. Other researchers have identified the problem that consensus assembly poses for the downstream analysis of variants and are moving towards alternative assembly approaches, such as graph-based assemblyGarrison et al. (2016).

### Haplotype recovery

The problem of haplotype recovery was first described by Lancia et al. (2001). Lancia introduced the first terminology and notation for “computational SNPology”^1^ (single nucleotide polymorphism) that served as a foundation for many other approaches and algorithms that followed. Lancia’s framework introduced the **SNP matrix** (typically denoted M): an *n* × *m* matrix encoding the binary allele (*A, B*) observed at each SNP site 1..*n* on each fragment 1..*m*. That is, *M*[*i*][*j*] is the allele observed at the *j*’th SNP of the i’th fragment (Lancia et al., 2001).

In this work Lancia introduced three optimisation problems:

- *Minimum fragment removal* (**MFR**);
- *Minimum SNP removal* (**MSR**); and
- *Longest haplotype reconstruction* (**LHR**).

All three approaches focus on removing the minimal number of rows or columns from the SNP matrix to correct “conflicts” until there exists a pair of haplotypes without conflicting evidence.

Lippert et al. (2002) introduced various solutions for the problems of MFR and MSR, but their main contribution was the definition of the *minimum error correction* (**MEC**) approach. MEC (also known as *minimum letter flip* (**MLF**)) attempts to find the minimum number of elements to “flip” in the SNP matrix M, such that a pair of haplotypes becomes feasible. Implementations of MFR, MSR and MEC/MLF include `Fast Hare, HapCUT` and others (Geraci, 2010; Lancia, 2016; Panconesi and Sozio, 2004; Bansal and Bafna, 2008).

All of these approaches focus on removing (or altering) observed evidence in the SNP matrix M until two compatible haplotypes can be defined. Each of these approaches has been demonstrated to be NP-hard. These methods typically fail to scale with the large sets of sequence data that have become common place since 2008 Aguiar and Istrail (2012).

Probabilistic approaches provide alternative methods that try to address this issue (Lancia, 2016; Li et al., 2004). Wang et al. (2006) introduced a polynomial time Markov chain based algorithm for the determination of diploid haplotypes. Markov models capture probabilities of transitions from one variant to the next, but have the memory-less property that each transition between the variant states is determined only by the state at the previous variant. Wang et *al*. tested increasing the “memory” (order) of the model to 3, however the results were not improved. This approach makes the assumption that the haplotypes to recover are from a diploid species, and that the genotypes of the allele pairs comprising the haplotypes are known in advance.

Aguiar and Istrail (2012) created `HapCompass`, which introduces a novel data structure: the *compass graph*, in which haplotype phasings correspond to spanning trees, in order to scale to the data output from modern sequencing methods. They later expanded on HapCompass (Aguiar and Istrail, 2013) to produce the first haplotype recovery algorithm to operate on polyploid genomes. However, it requires the ploidy to be specified in advance (diploid is assumed otherwise), which is unknown for metagenomes.

Evidently, the problem of haplotyping is a rich area of research, and has received much focus since it was first introduced in 2001, but the majority of modern haplotyping recovery software systems are not designed for metagenomic applications.

Existing algorithms are designed for single-species haplotype segregation, usually diploid species such as human. Typically they make easily violated assumptions, for example: SNP sites are bi-allelic (Ahn and Vikalo, 2015). However, metagenomes consist of an unknown number of organisms and their SNP sites can feature more than two alleles.

Approaches like those first presented by Lancia (Lancia et al., 2001) focus on the removal of errors. Such an approach would prove problematic if applied to metagenomic data, where errors are not just assumed, but are indistinguishable from both intra- and inter-species variation in the metagenome. For the same reason minimum letter flip (MLF) can be seen as unsuitable as it alters the observed information.

This type of problem is suited to Markov model based approaches. Markov models are particularly helpful on data sets without prior information (in their case, data errors, but in our case, contaminants: we know nothing of the species present, nor their distributions). We have applied a Markov model based approach to attempt to address the problem of haplotypes from metagnomic data sets.

In this work we introduce:

1. a novel probabilistic pseudo-graph data structure, **Hansel**, designed to store and provide an interface to pairwise co-occurring SNP evidence from sequenced reads.
2. an algorithm, **Gretel**: designed to take advantage of the `Hansel` data structure to load and traverse `Hansel` graphs to recover possible haplotypes from a *metahaplome*.

## Results

### The metahaplome

We define the metahaplome as the set of haplotypes for any particular genomic region of interest within a metagenomic data set.

### Hansel and Gretel

We present `Hansel` and `Gretel`, a data structure and algorithm for the recovery of haplotypes from a metahaplome. Advantages of our method include:

- recovers haplotypes from metagenomic data
- does not need *a priori* knowledge of the number of haplotypes
- makes no assumptions about the distribution of alleles at any variant site
- does not need to distinguish between sequence error and variation
- uses all available evidence provided by the raw reads
- does not require any user-defined parameters

We provide open source implementations for the data structure API (`Hansel`) and the haplotype recovery algorithm (`Gretel`) at https://github.com/samstudio8/gretel.

### Synthetic metahaplomes

We evaluate `Hansel` and `Gretel` on simple simulated metahaplomes constructed as described in our Methods.

We quantify success by evaluating each recovered haplotype against each of the generated haplotypes known to exist in the metahaplome. For each input, its best corresponding output is defined as the haplotype found by `Gretel` that shares the highest sequence identity with the generated input (Figure). We report the average of the best identity percentages for each input/output haplotype pair as the quality metric for our approach on synthetic metahaplomes.

Our results in Table 1 are promising. Best recovery rates (the highest recovery rate seen across the 10 randomly generated metahaplomes for that combination of haplotype length and number) are 100% for all but two (99.0%, 99.6%) of all tested lengths of metahaplomes containing up to 10 input haplotypes. We are able to recover at least one (but typically more) input haplotypes in their entirety, even for sequences consisting of 250 SNPs.

**Table 1:** Results of the synthetic metahaplome tests. The metahaplomes contained a known number of haplotypes of fixed size. Reads were randomly generated to span between 2–5 SNPs, with an approximate coverage of 3–5x for each haplotype. Each cell details the lowest (top), average (bold) and highest (bottom) best recovery rates (Figure) as discovered by `Gretel` over 10 repeats.

| Num/Len | 10 | 25 | 50 | 100 | 250 |
| --- | --- | --- | --- | --- | --- |
| 3 | 60.0 | 76.0 | 58.0 | 60.0 | 50.8 |
|  | **96.33** | **97.07** | **89.87** | **82.37** | **76.53** |
|  | 100.0 | 100.0 | 100.0 | 100.0 | 100.0 |
| 4 | 80.0 | 60.0 | 56.0 | 43.0 | 40.4 |
|  | **95.75** | **92.70** | **83.45** | **79.38** | **70.77** |
|  | 100.0 | 100.0 | 100.0 | 100.0 | 100.0 |
| 5 | 60.0 | 64.0 | 50.0 | 42.0 | 37.6 |
|  | **94.20** | **90.08** | **81.96** | **69.28** | **64.37** |
|  | 100.0 | 100.0 | 100.0 | 99.0 | 100.0 |
| 10 | 50.0 | 44.0 | 40.0 | 34.0 | 32.4 |
|  | **87.20** | **78.44** | **66.40** | **60.62** | **49.94** |
|  | 100.0 | 100.0 | 100.0 | 100.0 | 99.6 |
| 15 | 40.0 | 44.0 | 36.0 | 32.0 | 30.4 |
|  | **82.73** | **69.73** | **58.21** | **49.83** | **42.37** |
|  | 100.0 | 100.0 | 98.0 | 87.0 | 72.4 |
| 25 | 40.0 | 36.0 | 34.0 | 32.0 | 29.2 |
|  | **73.92** | **60.75** | **50.55** | **43.94** | **37.90** |
|  | 100.0 | 100.0 | 88.0 | 68.0 | 58.8 |
| 50 | 40.0 | 36.0 | 34.0 | 31.0 | 28.8 |
|  | **66.96** | **53.46** | **45.17** | **39.75** | **34.73** |
|  | 100.0 | 84.0 | 68.0 | 62.0 | 46.8 |

It is not surprising to observe that recovery success rates reduce with increased haplotype length (*i.e*. number of SNPs) and/or number of input haplotypes. Despite this, we report high recovery rates for haplotypes from metahaplomes with 100 SNPs or more, for small numbers of haplotypes.

### Metahaplomes from real genes

To extend our validation to reads derived from real sequence, we experimented with two metahaplomes constructed from two sets of real genes: dihydrofolate reductase *DHFR*, and Aminoacyl tRNA synthase complex-interacting multifunctional protein 1 *AIMP1*. DHFR and AIMP1 were selected as a test to evaluate whether variation across highly conserved genes could be recovered by our algorithm. To test the limitations of our method, we also evaluated our framework against a densely populated metahaplome of highly variable influenza sequences.

Our Methods section provides insight into the creation of the metahaplomes used for evaluation. For DHFR, AIMP1 and FLU-A7, we report the percentage of correctly recovered SNPs for each input and its corresponding best recovered haplotype, as determined by BLAST.

### DHFR

Table 2 presents the percentage of correctly recovered SNPs on the best haplotype recovered for each of the input sequences. In general, the algorithm performs well, both *BC070280.1* and *XR_63Ą888.1* are recovered with few errors. The recovered haplotype for the latter has a 98.9% recovery rate, when recovered from reads of length 120, with 15x coverage.

**Table 2;.** Results summarising the percentage of SNPs correctly recovered by `Gretel` for each DHFR metahaplome constructed with synthetic reads of a given size and coverage (see columns). Each cell is the percentage of correctly recovered SNPs on the best recovered haplotype for the metahaplome column and input gene row.

|  |  |  |  |  |  |  |  |  |
| --- | --- | --- | --- | --- | --- | --- | --- | --- |
| Read Length | 60 | 60 | 75 | 75 | 90 | 90 | 120 | 120 |
| ≈ Per-hap. Cover (5) | 15x | 7.5x | 15x | 7.5x | 15x | 7.5x | 15x | 7.5x |
| Number of SNPs | 79 | 67 | 87 | 78 | 95 | 78 | 95 | 92 |
| Gene ID | % | % | % | % | % | % | % | % |
| BC070280.1 | 94 | 79 | 92 | 91 | 96 | 92 | 96 | 96 |
| XR_634888.1 | 97 | 85 | 92 | 94 | 96 | 92 | 99 | 97 |
| KJ913115.1 | 84 | 55 | 69 | 88 | 92 | 64 | 87 | 90 |
| AK232978.1 | 71 | 40 | 77 | 69 | 84 | 65 | 87 | 89 |
| XM_012113510.1 | 62 | 70 | 77 | 76 | 81 | 85 | 82 | 77 |

Clearly, `Gretel` has particular difficulty recovering the *AK232978.1* and *XM012113510.1* haplotypes, which both demonstrate lower identities with the pseudo-reference. We observe that reads that are more divergent from the pseudo-reference sequence are discarded by alignment operations, denying `Gretel` the pairwise evidence needed to recover these less similar sequences. Despite this, on average *XM012113510.1* is recovered at over 75% accuracy.

### AIMP1

Table 3 presents the percentage of SNPs that are correctly recovered for the best haplotype recovered for each of the AIMP1 genes that are represented by synthetic reads in the metahaplome. Recoveries were not possible for either of the 60bp read metahaplomes, nor the 75bp 7.5x coverage meta-haplome as there existed a pair of SNPs too far apart to be spanned by at least one read in the data set. Of the results where analysis was possible, recovery rates are very high. Three of the five input genes have average recovery rates of over 96%; that is, at least 96% of the (on average) 70 variants are recovered correctly and match the expected input haplotype. We again have trouble recovering the sequence exhibiting the highest deviation from the reference *XM_006778898.2*, scoring an average recovery of just under 60%.

**Table 3:** Results summarising the percentage of SNPs correctly recovered by `Gretel` for each AIMP1 metahaplome constructed with synthetic reads of a given size and coverage (see columns). Each cell is the percentage of correctly recovered SNPs on the best recovered haplotype for the metahap-lome column and input gene row. Rows are ordered by decreasing identity to the pseudo-reference.

|  |  |  |  |  |  |  |  |  |
| --- | --- | --- | --- | --- | --- | --- | --- | --- |
| Read Length | 60 | 60 | 75 | 75 | 90 | 90 | 120 | 120 |
| ≈ Per-hap. Cover (5) | 15x | 7.5x | 15x | 7.5x | 15x | 7.5x | 15x | 7.5x |
| Number of SNPs |  |  | 68 | 62 | 74 | 72 | 71 | 76 |
| Gene ID |  |  | % |  | % | % | % | % |
| NM_004757.3 | - | - | 97 | - | 96 | 96 | 96 | 99 |
| XM_003829985.2 | - | - | 96 | - | 97 | 97 | 96 | 96 |
| XM_001170763.4 | - | - | 96 | - | 97 | 97 | 97 | 99 |
| XM_013365392.1 | - | - | 94 | - | 93 | 88 | 97 | 89 |
| XM_006778898.2 | - | - | 60 | - | 49 | 51 | 68 | 70 |

### FLU-A7 (metahaplome of 71 haplotypes)

We test the limitations of our approach with a metahaplome of 71 highly variable Influenza A Segment 7 haplotypes. For brevity we summarise in Table 4 the recoveries obtained across all 71 haplotypes and describe the percentage of SNPs recovered for the worst, average and best recovered haplotypes. Particularly impressive is the best haplotype recovered from the 120bp 7.5x data set, where 99% of the 264 SNPs agreed with those on its corresponding input gene. On average, the haplotype with the lowest sequence identity to the pseudo-reference has an worst recovery rate of 52.5%, which is far better than expected by chance alone.

**Table 4:** Results summarising the percentage of SNPs correctly recovered by Gretel for the FLU-A7 metahaplome (71 highly variable Influenza A Segment 7 haplotypes) constructed using synthetic reads of different sizes and coverages (see columns). Cells show the percentages of correctly recovered SNPs for the best recovered haplotype, average haplotype and worst recovered haplotype.

| Read Length<br>≈ Per-hap. Cover (71)<br>#SNPs | 60<br>15x<br>241 | 60<br>7.5x<br>227 | 75<br>15x<br>256 | 75<br>7.5x<br>236 | 90<br>15x<br>263 | 90<br>7.5x<br>242 | 120<br>15x<br>289 | 120<br>7.5x<br>264 |
| --- | --- | --- | --- | --- | --- | --- | --- | --- |
|  | % | % | % | % | % | % | % | % |
| Best | 94 | 88 | 96 | 94 | 96 | 95 | 95 | 99 |
| Average | 69 | 66 | 71 | 68 | 72 | 68 | 75 | 72 |
| Worst | 50 | 45 | 54 | 50 | 53 | 48 | 63 | 56 |

It should be noted that for each of these metahaplomes, between 25% and 50% of the generated synthetic reads were discarded. It is highly likely with finer tuning, a different alignment algorithm, or an alternative approach to alignment, we could drastically improve the already reasonable scores presented here.

## Discussion

We have provided a description of a previously undefined problem and made advances in the recovery of haplotypes from a metagenome. We offer the term metahaplome to represent the set of haplotypes for any particular region of interest within a metagenomic data set.

We introduced Hansel, a novel data structure which encodes the variation seen across a **metahaplome**. Hansel permits traversal of that variation like a graph, yet allowing for probabilistically weighted edges to consider the state of the haplotype recovered so far. We also introduced *Gretel*, an algorithm capable of traversing the metahaplome represented by *Hansel* for the recovery of genuine haplotypes from a metahaplome constructed from the raw reads of a metagenomic data set. For the first time, it is possible to computationally extract variants of commercially relevant genes.

Together `Hansel` and `Gretel` form a new framework for the recovery of haplotypes in metagenomes, allowing data sets where short read length has previously restricted effective analysis.

### Performance and tractability

Without an annotated metagenome, it is clearly difficult to quantify the effectiveness of our approach. The testing presented here has been performed with data simulated from real genes in order to explore the limitations of the algorithm.

As described in our methods, initial testing was completed with randomly generated haplotypes to measure performance with regard to both haplotype length and number of haplotypes. We demonstrate very high recovery rates, even in the presence of many SNPs.

We evaluated the approach with synthetic reads generated from metahaplomes consisting of mixtures of real genes (DHFR and AIMP1) and demonstrated it is possible to recover haplotypes from short read data accurately. Successful recoveries can be made with our framework even in the presence of many haplotypes. We demonstrate high recovery rates in the FLU-A7 metahaplome, containing short reads generated from 71 highly variable influenza sequences.

However, our results have also demonstrated that our approach is strongly biased by the alignment of reads against the pseudo-reference. During testing of both the DHFR and AIMP1 data set, it was found that many synthetic reads would not align back to the pseudo-reference. Input genes *(i.e*. haplotypes) for the DHFR and AIMP1 metahaplomes were selected with decreasing sequence identity from the pseudo-reference (the pseudo-reference). Unfortunately, when short reads were generated, it was found that `bowtie2` discarded up to 20% of the synthetic reads, particularly those yielded from less similar sequences.

This is of course not unexpected: it should not be a surprise that sequences with lower identity to the target are likely to eventually fall below some threshold and be discarded. However, this does raise an important caveat to our work: both assemblers and aligners will exert influence over the tractability of how many and how accurately haplotypes in a given metahaplome can be recovered. Many software packages that perform these tasks have expectations and make assumptions that are not ideal in the case of recovering haplotypes from a metagenome. Here, the discarding of reads denies `Hansel` and `Gretel` access to critical evidence require to reconstruct those particular haplotypes.

It should be noted that the pseudo-reference is not used by `Hansel` or `Gretel`, it serves only as a common sequence on which to align raw reads before calling SNPs. Very high recovery rates on sequences that share identity with the pseudo-reference are a reflection of the strength of our approach, and not a trivial recovery.

Perhaps most significantly, the tractability of the problem is bound by the quality of the data available. As stated by Lancia in 2001, it is entirely possible that, even without error, there are scenarios where data is insufficient to successfully recover haplotypes and the problem is rendered impossible Lancia et al. (2001). Indeed, as explained in our Methods section, there are multiple potential inconsistencies that can occur in the alignment that are not trivial to address.

It should be noted that although our framework has been designed with the recovery of haplotypes in a metagenome at a gene level *(i.e*. variants of a gene involved in a catalytic reaction of interest, such as degradation of biomass) in mind, given sufficient coverage of SNPs, our approach could work on regions significantly longer than that of a gene.

Regarding time and resource requirements, Hansel and Gretel is designed to work on all reads from a metagenome that align to some region of interest on the pseudo-reference. Typically these subsets are small (on the order of 10–100K reads) and so our framework can be run on an ordinary desktop in minutes, without significant demands on disk, memory or CPU.

### Advantages of our approach

In contrast to other methods, our framework aims to make as few assumptions as possible. `Gretel` is designed for metagenomic data sets where the number of haplotypes is unknown. Whilst `HapCompass` is designed to identify haplotypes for a polyploid organism, it requires advance knowledge of the number of expected haplotypes Aguiar (2014) which is unknown for metagenomic data sets.

Most SNP calling algorithms discard SNP sites that feature three or more alleles *(i.e*. non bi-allelic sites) as errors, or under the assumption that input data represents sequenced reads from a diploid species Ahn and Vikalo (2015). Although `ParticleHap` (Ahn and Vikalo, 2015) relaxes this assumption by incorporating genotype calling, it is to reduce the risk of erroneously called genotypes preventing reconstruction of the two haplotypes for a diploid genome.

Many existing methods rely on discarding or altering observed SNPs until a pair of haplotypes can be determined. `Hansel` and `Gretel` uses all available pairwise observations and works to recover the most likely haplotypes. Unlike other methods, we do not assume that the observed evidence must be contaminant or sequencing error that needs discarding or altering to recover the real haplotypes. Although sequence error is an inevitability, errors will be poorly supported by read data and are unlikely to form components of recovered haplotypes.

## Future work

`Gretel` is a proof of concept. Although we have demonstrated success in our Results, there are still other sources of evidence not currently used by our algorithm — namely paired end reads and alignment base quality scores. Read pairs will certainly provide useful co-occurrence information for SNPs that span some known insert, however careful consideration on how to integrate this data is necessary. The order of the Markov chain that constructs haplotypes will typically be small, as it represents the number of selected variants from the current head of a path to include when considering probabilities for which variant should come next. As reads typically span only a few SNP sites, it is not effective (and can be detrimental) to set the “lookback” parameter *L* to a value high enough to consider variants seen at the other side of an insert.

Additionally, the second read of a pair will provide evidence for variants that are likely appear after an insert (given the variants seen in the first read of the pair), but no evidence for what variants should be selected during the insert. During recovery, this potentially puts us in the position of having evidence for what a variant a few positions ahead of our current location should be, but no idea of how to get there. A solution may be to permit `Gretel` to fill in future variants given paired end evidence (if available) as placeholders and backpropagate towards the head of the path to predict which variants are likely to appear given the placeholder observed in the future. This is possible as the pairwise information stored in `Hansel` by `Gretel` is not directional. Observations are bidirectional, and although the structure presented by `Hansel` is a directed graph, the direction can be either forwards or backwards, just not both. This would allow reconstruction of haplotypes from either end of a gene.

Alignment scores would permit some form of weighting mechanism to be applied to observations. Low confidence base calls can be considered as less informative than those with high calling confidence, but experimentation on how to reliably tune this parameter for pairwise information (how should we weight a pair of variants where one call is good, and the other is bad) would be needed.

Our largest obstacle is that of *smoothing*. In general, smoothing tries to reduce overfitting of a model. Here, we want to avoid scenarios where SNP sites with very low read coverage (and thus few informative observations) are assumed to be fully representative of the true variation. Many of the alignment artefacts described in our Methods section (Figure 2) as posing a problem for reconstruction *(e.g*. SNP pairs with few valuable observations, SNP sites that are not connected by reads) can begin to be better addressed with an appropriate model for smoothing. Future work aims to build upon the simple add-one smoothing found in our current model to incorporate more intelligent smoothing of low frequency observations.

**Figure 1:**
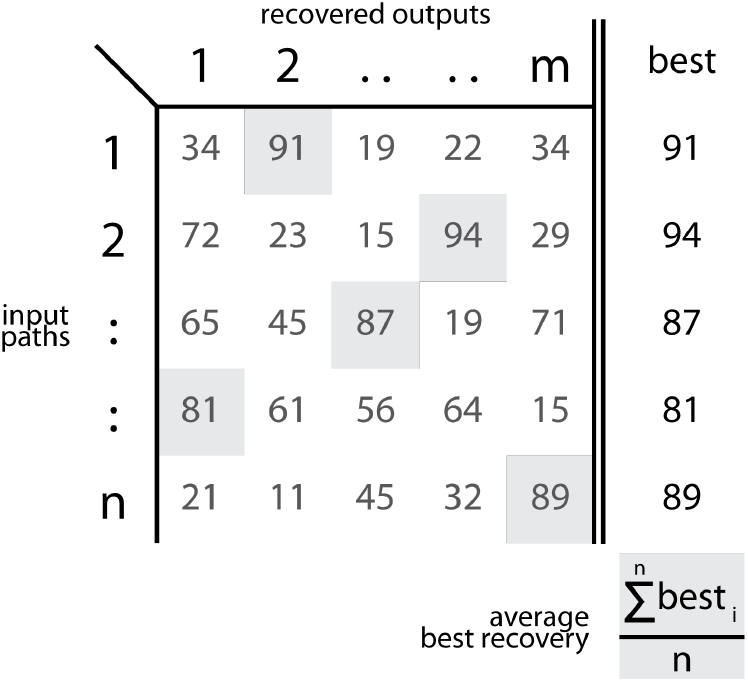
An example demonstrating the calculation of the recovery rate of haplotypes from generated synthetic metahaplomes. Rows represent known input haplotypes, columns represent the haplotypes recovered by `Gretel`. Cells report the percentage sequence identity of that input-output pair. The average best recovery is the sum of the best identity percentage for each input haplotype, divided by the number of inputs. Values are for illustrative purposes only. Table 1 reports the average scores of the best recovery rate for each generated haplotype.

**Figure 2:**
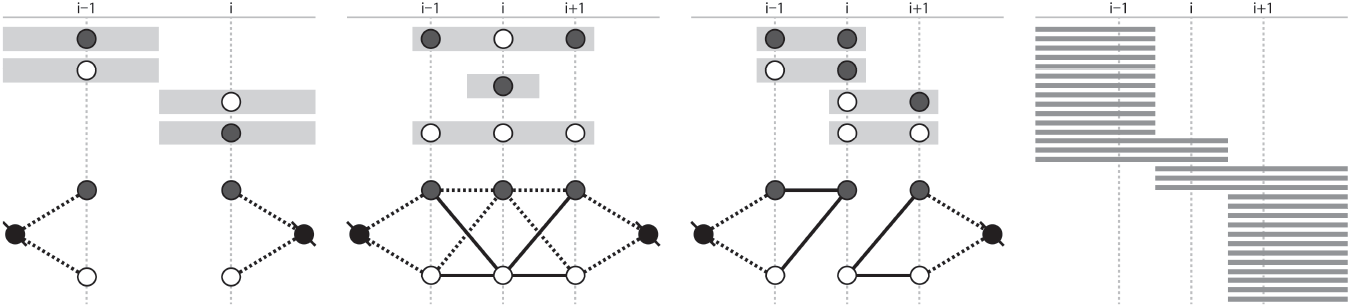
Artefacts of read alignments that cause difficulties for recovery of haplotypes from metahaplomes (a) Reads do not span a pair of SNPs, no pairwise evidence describing how the graph can be traversed is available (b) single SNP reads do not provide sufficient evidence about their source or destination variants and add many potential edges to the graph (c) coverage of reads exists between a pair of SNPs but the information available is insufficient to determine paths through the graph (d) low coverage between SNP sites can cause decisions to be biased

`Hansel` and `Gretel` only consider SNPs. It is likely that real haplotypes will exhibit insertions and deletions (indels) against a pseudo-reference. Further thought and experimentation is needed to devise a methodology that permits the consideration of indels by our approach. Whilst an initial solution may be to add an additional symbol to represent an indel to the `Hansel` structure, we must find a way to incorporate this evidence into a structure that currently only considers paired SNP variation.

`Gretel’s` algorithm involves a greedy bias. We assume the “best” haplotype is the most likely haplotype, and that it can recovered by selecting the edge with the highest probability at each SNP. However there are likely to exist solutions whose overall likelihood may be higher if we permitted `Gretel` to look ahead along the path to inspect a small number of future choices. It is possible that this will alter little in in practice, as each read only spans a small number of SNPs.

The “lookback” parameter *L* does have some influence on both the efficiency and accuracy of the recovery. A meaningful choice for this parameter is to use the mean number of SNPs covered per read.

As briefly described in our Methods, Gretel will continue to generate possible haplotypes until the available evidence in Hansel is exhausted. Thus our algorithm is capable of determining its own stopping criteria. Recovered haplotypes can be ranked according to metadata provided by Gretel such as the log likelihood of that haplotype occurring given the variation observed across the original raw reads. We intend to explore additional metrics that may be used to determine other methods for scoring (and filtering) returned haplotypes.

Finally, we would like to investigate potential methodologies for abandoning the requirement of a common reference (that is, in our terminology, the assembly, or pseudo-reference) and working solely with read data. A metagenomic assembly provides both a convenient proxy for the raw reads (that is, a gene found or predicted on the pseudo-reference can be assumed to have some affinity with reads that align to the same location), and a pseudo-reference to align reads against and call for SNPs. However, the processes of assembly, alignment and SNP calling each make assumptions and decisions that simplify or discard data, reducing the evidence available for the accurate recovery of haplotypes. With fast and efficient sequence search alternatives(Buchfink et al., 2015) it is easier to conduct sequence similarity searches across very large sets of metagenomic reads. Literature also exists for reference-free SNP callingIqbal et al. (2012), which both offer opportunities for introducing fewer assumptions and maintaining the in-tegrity of variants observed across metagenomic data before they reach a framework such as `Hansel` and `Gretel` for the recovery of haplotypes, from a metahaplome.

## Conclusion

In this work we have introduced a definition for the **metahaplome**. We provide both a definition and implementation of a novel data structure, and a proof of concept algorithm, that together represent a framework for the reconstruction of haplotypes from metagenomic data sets. We demonstrated promising recovery rates and described many interesting avenues for future work and analysis.

We aim to empirically prove that `Gretel` is capable of reconstructing real haplotypes by using its output to design primers to extract a commercially interesting enzyme and its variants from a metagenome and confirm the results using single molecule sequencing.

For the first time, we have demonstrated computational techniques that are capable of using metagenomic reads to reconstruct viable proteins responsible for interesting catalytic reactions in a microbial community; a task that existing computational methods and alternative laboratory techniques such as rational design have struggled to achieve.

Our work is an advance in computational methods for extracting exciting exploitable enzymes from metagenomes.

## Methods

### The metahaplome

To enable recovery of haplotypes from a metahaplome for a metagenomic data set, we assume the following is available:

- A set of *raw reads* from a metagenome
- An assembly of those reads (the *pseudo-reference*)
- A region of interest on the assembly (the *target*)
- An *alignment* of the raw reads, against the pseudo-reference
- A list of *positions* at which single nucleotide variations occur over the aligned reads

A pseudo-reference can be generated by assembling sequenced reads known to come from a metagenomic data set, with an assembler such as `VelvetZerbino` and Birney (2008). Selecting a suitable assembly algorithm and the dark art of choosing parameters such as k-mer size is left as an exercise to the reader. A region of interest on the assembly may be identified by homology search or gene prediction. Raw reads are filtered by whether or not they “map back” to the target region according to an alignment tool such as `bowtie2`Langmead and S. (2012). Reads that fall outside the target of interest (*i.e*. reads that do not cover any of the genomic positions covered by the target) are discarded. Variation at single nucleotide positions across reads along the target, can then be called with a SNP calling algorithm such as that provided by `samtools`Li et al. (2009) or `GATK`DePristo et al. (2011). Alternatively, one may determine any position that features at least one (or some number of) reads that disagree with one-another or the pseudoreference as a SNP.

The combination of aligned reads, and the locations of single nucleotide variation on those reads can be exploited to recover real haplotypes from the metahaplome: the collection of haplotypes that exist for the given region of interest on the metagenome.

Our ability to recover haplotypes from a metahaplome depends on the quality and coverage of the available reads and their alignment. Figure 2 depicts several inconsistencies in aligned reads that make recovery of haplotypes more difficult.

### Hansel: A novel data structure

We present **Hansel**, a probabilistically-weighted, graph-inspired, novel data structure. Although the structure can be traversed like a graph, its underlying representation is a four dimensional array whose elements represent the number of observations of a given symbol *A* appearing at position *i*, co-occurring with symbol *B* at position *j*.

This representation differs from the typical SNP matrix model (Lancia et al., 2001) that forms the basis of many of the surveyed approaches. Rather than a matrix of SNP columns and fragment rows, we discard the concept of a fragment entirely and aggregate the evidence seen across all fragments by position.

For each possible pairing of symbols (*i.e*. AA, AC,… TG, TT), the `Hansel` structure keeps a matrix, whose elements record the number of observations of those symbols co-occurring on a read. More specifically, an element in the `Hansel` matrix *H, H*[*A,B,i,j*] stores the number of times symbol *A* at position *i* has been seen to co-occur with symbol *B* at position *j*. For example, the number of times a `C` at SNP position `3`, appears on the same read as a `T` at SNP position `6`.

Although this structure may appear limited, the data can be exploited to build other structures. For example, if one considers *H*[*A, B*, 1,2] for all possible *A* and *B*, one may list the available options for transitioning from position 1, to position 2. Extending this to consider every element *H*[*A, B,i,i* + 1], for all possible combinations of symbols *A* and *B*, and all SNP positions *i*, we can construct a SNP transition graph *G* (Figure 3).

**Figure 3:**
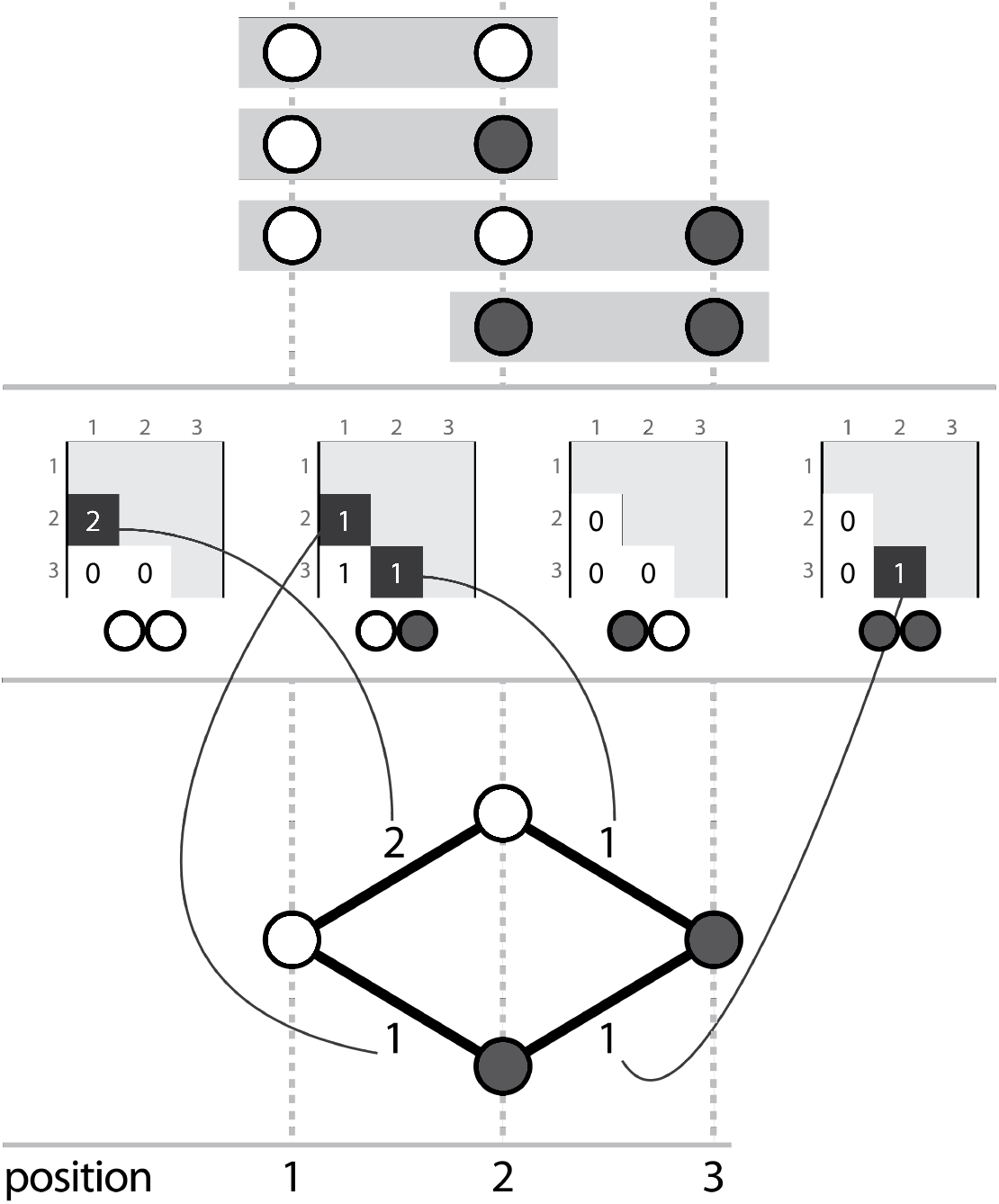
Three corresponding representations, (a) aligned reads, (b) the Hansel structure, (c) a graph that can be derived from the Hansel structure

Intuitively, one may traverse *G* by selecting edges with the highest weight (where edge weights may be defined as the number of reads in which the current node was observed to occur with a given next node) to recover chains of symbols that represent an ordered sequence of SNPs that constitute a haplotype.

However, the data cannot be fully represented with a graph such as that seen in Figure 3 alone. This representation defines a constraint whereby edges may only join adjacent SNPs and so cannot encode any information as to which non-adjacent polymorphisms co-occur. Without considering information about non-adjacent SNPs, one can traverse the graph to create paths that don’t exist in nature (Figure 4).

**Figure 4:**
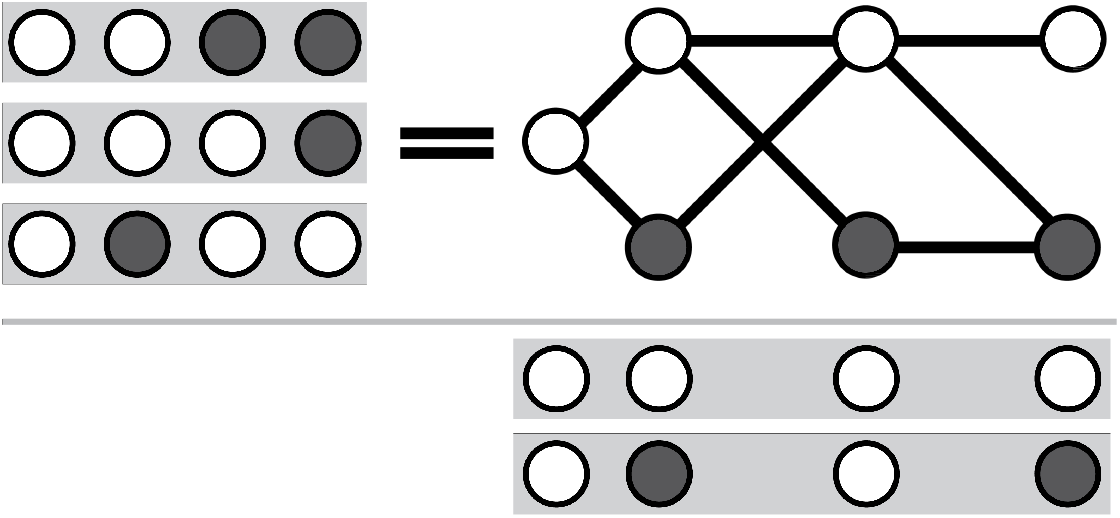
Considering only adjacent SNPs, one may create paths for which there was no actual observed evidence. Here, the reads {0011, 0001, 0100} do not support either of the results {0000, 0101}, but both are valid paths through a graph that permits edges between pairs of adjacent SNPs.

To recover real haplotypes accurately, we must consider more than just the head of the path. The `Hansel` structure is designed to store pairwise co-occurrences of all SNPs (not just those that are adjacent), as seen across all reads. Thus we can weight edges in the graph based not just on the number of reads containing the current node, and a possible next node, but previously selected nodes too.

However, this does come at a cost. We describe `Hansel` as “graphinspired”, as allowing edge weights to depend on more than just the current node (that is, allowing an edge to be weighted with information from the `Hansel` structure that does not pertain to that edge specifically) leads to several differences between the `Hansel` structure, and a weighted directed acyclic graph. Whilst these differences are not necessarily disadvantageous, they do change what we can infer about the structure.

Now, the structure of the graph is effectively unknown in advance (Figure 5). That is, not only are the weights of the edges not known ahead of traversal, but the entire layout of nodes and edges is also unknown until the graph is explored (although, arguably this would be true of very large simple graphs too). Indeed, this means it is also unknown whether or not the graph can even be successfully traversed.

**Figure 5:**
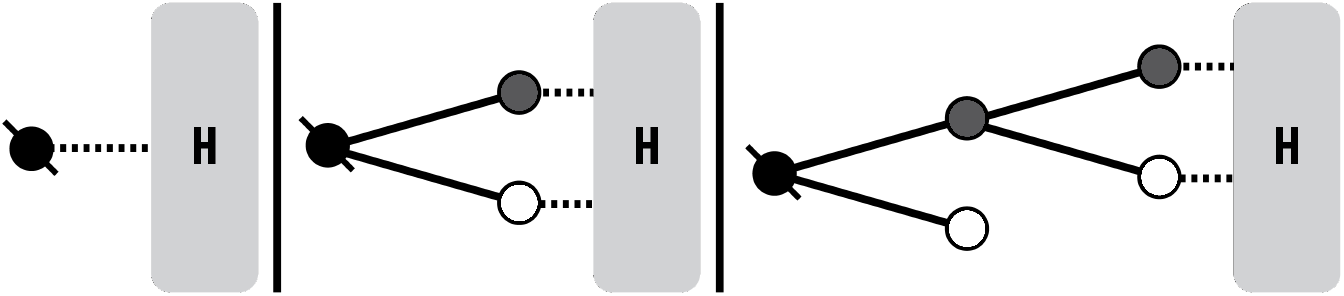
The physical characteristics of the graph are unknown *a priori*. The metahaplome must be explored to determine its components. Only branches that are selected are explored further.

Secondly, the graph is dynamically weighted. The current path represents a memory that affects the availability and weights of outgoing edges at the current head node. Edge weights are calculated probabilistically *during* traversal and depend on both the distribution of variants observed at the position to traverse to next, but also some number of the variants that have been encountered and selected thus far in the path. Please refer to our supplementary materials for information on how probabilistic weightings are calculated.

**Figure 6:**
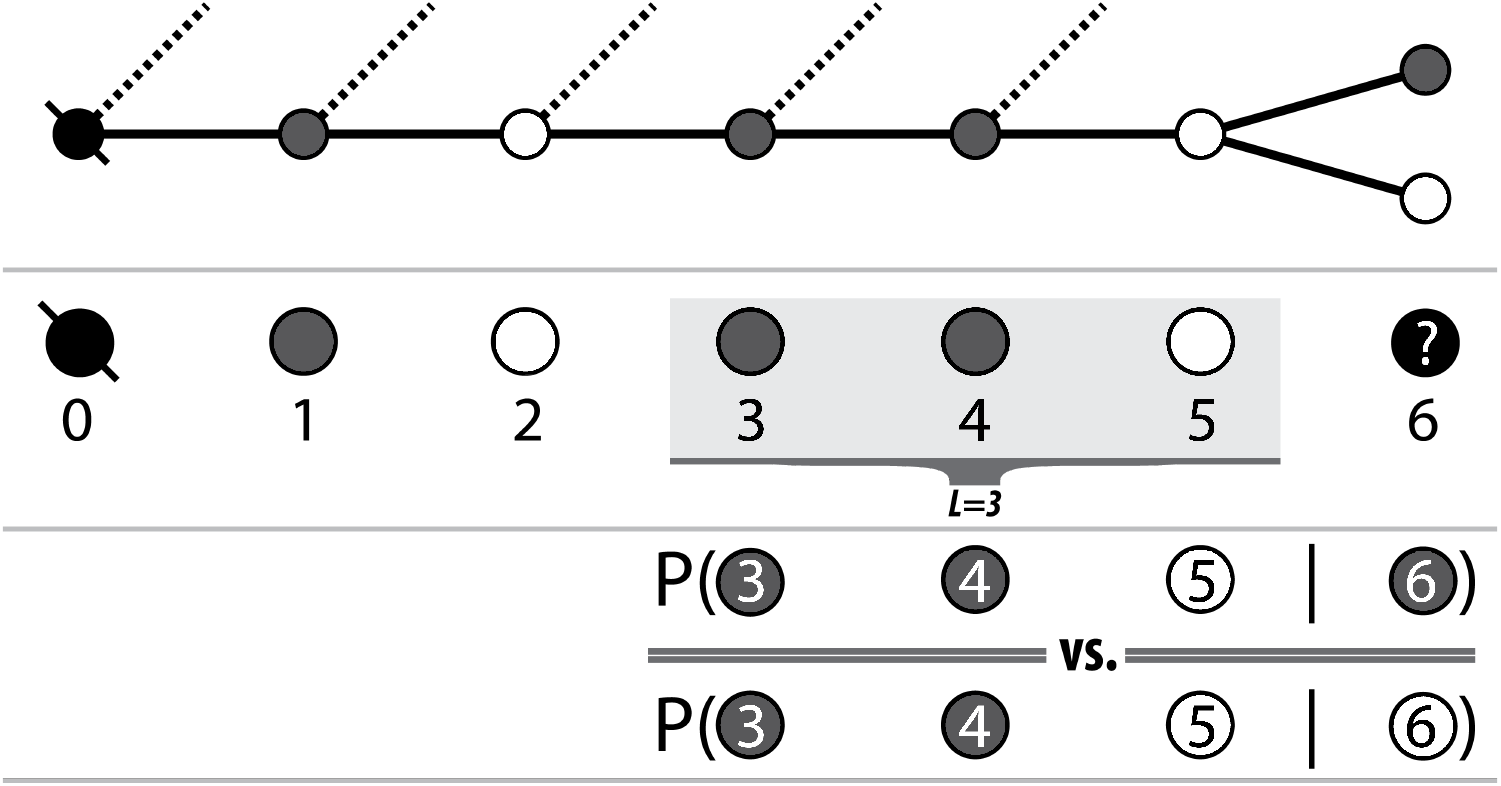
Pairwise conditionals between L last variants on the observed path, and each of the possible next variants are calculated and the best option (highest likelihood) is chosen

Effectively, a “fog of war” exists over the graph that is only dissipated as it is traversed. In exchange for these minor caveats, we have a data structure that permits graph-like traversal that is intrinsic to our problem definition, whilst utilising informative pairwise SNP information collected from observations on raw metagenomic reads. `Hansel` fuses the advantages of a graph’s simple representation (and its inherent traversability) with the advantage of a matrix of itemsets (that permit storage of all pertinent information).

We provide an open source implementation of `Hansel` in the form of a Python package that exposes a friendly interface capable of managing and querying pairwise observations to a graph-inspired data structure for determining likely chains of sequences from breadcrumbs of evidence.

### Gretel: An algorithm for recovering haplotypes from metagenomes

We introduce `Gretel`, a proof of concept algorithm designed to interface with the `Hansel` data structure to recover the most likely haplotypes from a metahaplome. To obtain likely haplotypes, `Gretel` traverses the probabilistic graph structure provided by `Hansel`, selecting the most likely SNPs at each possible node (*i.e*. traversing edges with the greatest probability), given some subset of the most recently selected nodes in the path so far. At each node, a Markov chain of some order L is employed to predict which of the possible variants for the next SNP is most likely, given the last L variants in the current path.

Execution of `Gretel` can be broken into the following steps:

1. Parse alignments and construct “SNP strings”
2. Populate `Hansel` structure with pairwise observations
3. Exploit the `Hansel` graph API to incrementally recover a path until a variant has been selected for each SNP position

- Query for the available transitions from the current head node to the next SNP
- Calculate the probabilities of each of the potential next variants appearing in the path given the last *L* variants
- Append the most likely variant to the path and traverse the edge
4. Re-weight used observations in `Hansel` to allow for new path
5. Repeat (3–4) until the graph can no longer be traversed or an optional additional stopping criterion has been reached

Haplotypes are reconstructed as a path through the `Hansel` structure, one SNP at a time, linearly, from the beginning of the sequence. At each SNP position, the `Hansel` structure is queried for the variants that were observed on the raw reads at the next position. `Hansel` also calculates the conditional probabilities of each of those variants appearing as the next SNP in the sequence, using a Markov chain that makes its predictions given the current state of the observations in the `Hansel` matrix and the last *L* selected SNPs. `Gretel’s` approach is greedy: we only consider the probabilities of the next variant. Our razor is to assume that the best haplotypes are those that can be constructed by selecting the most likely edges at every opportunity.

Once a path is completed (a variant has been chosen for all SNP sites), the observations in the `Hansel` matrix are re-weighted by `Gretel`. Whilst our framework is probabilistic, it is not stochastic. Given the same `Hansel` structure and operating parameters, `Gretel` will behave in a deterministic fashion and return the same set of paths every time. However we are interested in recovering multiple, real haplotypes from a metahaplome, not just one. `Hansel` exposes a function in its interface for the re-weighting of observations. Currently, `Gretel` reduces the weight of each pairwise observation that forms a component of the completed path - in an attempt to erase evidence for that haplotype existing in the metahaplome at all, allowing evidence for other paths to now direct the probabilistic search strategy.

Finally, `Gretel` outputs recovered sequences as FASTA, requiring no special parsing, or munging of results to be able to conduct further sequence analyses.

### Testing methodology

Our testing evaluates the performance of our framework against metahaplomes consisting of synthetic reads derived from both randomly generated haplotypes, and also haplotypes created from real gene sequences.

There are no metagenomic data sets with rich haplotype level annotations. We chose to use synthetic data sets to evaluate our framework under known, controllable conditions, and to afford us the ability to actually quantify the accuracy of recovered haplotypes from a metahaplome.

In this section, we provide an overview of the methods to generate metahaplomes for both random haplotypes, and haplotypes based on real genes. We describe our approach for evaluation of our work. In our Results, we demonstrate the effectiveness and limitations of our framework.

### Synthetic metahaplomes

With the desire to first test our approach on data that was as simple as possible, we generated small, synthetic metahaplomes which contain a known number of randomly generated haplotypes, each of the same fixed length. Every position on the haplotype is regarded as a site of variation. Each of the random haplotypes in the metahaplome is permitted to select any of the four base pairs at random, for each genomic position. An arbitrary haplotype is drawn from the metahaplome and chosen to be the ‘pseudoreference’. We generate short reads from the other remaining haplotype sequences, each read is between 3–5bp (thus, 3–5 SNPs). These reads are constructed by sliding a window along each haplotype, whilst also varying the size, and overlap of those windows in an attempt to introduce some element of realism to the data.

As we know the location and width of each such window, we can append synthetic alignments to a SAM file without having to invoke an actual aligner. This is particularly important, given that the randomly generated haplotypes represent strings of SNPs and exhibit low sequence identity between one another. Combined with their very short nature, this prevents alignment tools from assisting us with generating an alignment format for input to our algorithm on such data. The majority of sites are tri- or tetra-alleleic and so for the same reason, the VCF is produced by our metahaplome generator, rather than an established diploid-assuming SNP caller. Both a SAM alignment, and a VCF are generated by our tool, circumventing the assumptions and biases of real aligners and SNP callers.

We detail the parameters and options of our metahaplome generator in the supplementary materials. The code is open source and freely available via our data and testing repository, online: https://github.com/samstudio8/gretel-test

### DHFR and AIMP1

We chose an arbitrary DHFR and AIMP1 gene from GenBank to serve as the ‘master’ sequence (*i.e*. the pseudo-reference) for their respective metahaplomes.

A discontinuous megaBLAST was conducted for both of the DHFR and AIMP1 masters. From each, a set of five related but arbitrary genes of decreasing sequence identity (DHFR:≈ 99.8%, 97%, 93%, 90%, 83% and AIMP1: ≈ 99.9%, 99.6%, 99.3%, 92%, 91%) were selected.

Each of the five genes were broken into reads with a uniform size and overlap. Resulting reads were aligned back to the master with `bowtie2`. Variants were called on the alignment with a script that determined all non-uniallelic locations as SNPs. The DHFR data yielded between 65–90 SNPs per 564bp metahaplome, the AIMP1 data yielded 60–75 SNPs per 939bp metahaplome.

For testing, multiple DHFR and AIMP1 metahaplomes were generated. The read size was uniform and haplotype recovery was measured for metahaplomes populated with reads of size 60, 75, 90 and 120. Per-haplotype coverage was estimated by dividing the average coverage observed by `samtools depth`, by the number of input haplotypes. Per-haplotype coverage was adjusted by increasing or decreasing the overlap of the reads.

Quantification of recovery rate proved much more difficult for data yielded from discontinuous searches. The current implementations of `Hansel` and `Gretel` is gap agnostic. Reads with insertions or deletions are handled by parsing their BAM CIGAR strings and adjusting the nucleotide that appears at the SNP sites that follow accordingly. However, although with this method the SNP is technically correct, information about the size and location of the gap is lost. Thus when the recovered haplotypes are written to FASTA, they are gapless. Testing input and output haplotypes base-for-base is therefore not possible, any gap in the input haplotype will be ignored and `Gretel`’s performance will be reported poorly, even if the SNPs themselves are all correct (but in the wrong position).

We define the haplotype recovery rate from the DHFR and AIMP1 metahaplomes by BLAST. The known input genes are used as queries against a BLAST database constructed from the newly recovered output haplotypes from `Gretel`. For each input gene, the best output haplotype is defined as the path with the best BLAST hit, determined by bit score. In our Results, we report the proportion of SNPs that are correctly recovered on each gene’s best output haplotype.

### FLU-A7

A data set of 772 Influenza A (Segment 7) sequences were obtained from GenBank, requesting any sequences deposited from July 1st 2016 to July 25th 2016. After removing a majority of duplicate sequences, 72 sequences, of mean length 1009bp remained. One sequence was randomly selected as our pseudo-assembly. The remaining 71 sequences were aligned to the master and sharded into synthetic reads via the same method as described in our Methods for the DHFR and AIMP1 metahaplomes. SNPs were called in the same way, determining any site without a unanimous consensus as a variant site. Regardless of parameters provided to the read generator, there were typically at least 250 SNPs observed over the 982bp metahaplome. We designate the data set *FLU-A7*.

### Ranking haplotypes recovered from a metagenome

Of course, with knowledge of the input haplotypes that we expect to recover, we are able to quantify our approach. For real metahaplomes, we need a mechanism to differentiate false positives, or rank our confidence in the returned haplotypes.

Future work will investigate this in more depth, but currently, in addition to the sequences themselves, `Gretel` outputs a ‘crumbs’ file — a whimsical name for a simple, tab delimited format — contains metadata for each of the recovered sequences: log probability of that sequence existing given the evidence seen overall, how much of the evidence in `Hansel` that particular sequence was supported by, and how much of that evidence was re-weighted as a result of that path being chosen.

Currently, `Gretel` will continuously recover paths out of the remaining evidence until it encounters a node from which there is no evidence that can inform the next decision.

## Data Access

Our Hansel and Gretel framework is freely available, open source software available online at https://github.com/samstudio8/hansel and https://github.com/samstudio8/gretel, respectively.

The scripts used to generate metahaplomes and synthetic reads for both the randomly generated and real-gene haplotypes, and the testing data used to evaluate our methods is also available online via https://github.com/samstudio8/gretel-test

## Acknowledgments

CJC was funded by the Biotechnology and Biological Sciences Research Council (BBSRC) Institute Strategic Programme Grant, Rumen Systems Biology, (BB/E/W/10964A01). WA is funded through the Coleg Cymraeg Cenedlaethol Academic staffing scheme. SN is funded via the Aberystwyth University Doctoral Career Development Scholarship and the IBERS Doctoral Programme.

## 1 Supplementary Materials

### 1.1 Hansel as a graph

Consider an alphabet of symbols, Σ (*e.g.* {*A, C, G, T*}) and a list of *m* SNP positions 1..*m*. As described in our article, the `Hansel` structure *H* can be considered as a graph *G* = (*V, E*). Here, we define *V*, and *E*:

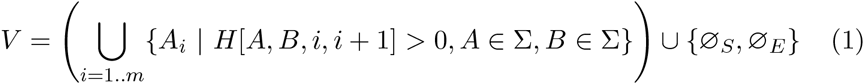

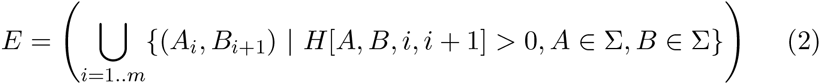

*V*, the set of nodes (or vertices), containing an element for all pairings of a symbol in the alphabet, and position where at least one read contains that symbol at *i*, and has coverage for at least *i* + 1.

*E*, the set of edges, where an edge (*A_i_, B*_*i*+1_) is determined to exist in *E* if there exists at least one read whereby symbol *A* was observed at position *i* to co-occur with symbol *B* at SNP position *i* + 1.

It should be noted, that although *G* can be constructed from *H* such that it is undirected and contains cycles, both properties lead to nonsensical haplotypes. Under such circumstances, `Gretel` could construct a path that visits multiple nodes that appear at the same *i*, or a trail that visits the same node multiple times. Such sequences would be meaningless in the context of haplotype construction, thus the interface to `Hansel` acts in such a way, that *G* is a directed, acyclic graph.

We can define a haplotype as an alternating sequence of nodes (*υ* ∈ *V*) and edges (*e* ∈ *E*). A path must always start and end at the special sentinel symbols *∅_S_* and *∅_E_*, respectively.

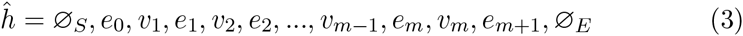

Although, as only one directed edge between some *υ_i_* and *υ*_*i*+1_ may exist, we can define 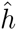 simply as a sequence of *υ* ∈ *V*:

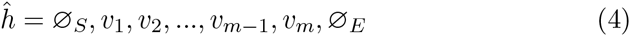

### 1.2 Probabilistic edge weights

However, as we described in our paper if the construction of *G* does not consider elements in *H*[*A,B,i,j*] where *abs*(*i − j*) > 1 it is likely one will recover haplotypes that do not actually exist.

Given the pairwise information available in *H*, for both adjacent, and non-adjacent SNPs, across all reads, we described that edges in the graph *G* derived from *H* can be weighted probabilistically.

We attempt to determine the next most likely symbol in a sequence, considering both the marginal distribution of symbols at the next position and the likelihood of those symbols appearing next, given an already observed partial sequence into account.

That is, the next symbol *υ*_*i*+1_ in a path depends not only on the current symbol (*υ_i_*) but some number of previous symbols (*υ_i−1_,υ_i−2_…υ_0_*).

The outgoing edges from *υ_i_* are probabilistically weighted by exploiting the observations stored in the `Hansel` structure to create probabilities, that then determine the likelihood of moving from some *υ_i_* to each of the possible *υ*_*i*+1_.

We take a Bayesian approach to the problem of probabilistically weighting edges in `Hansel`’s graph representation. We define the probability of selecting *υ*_*i*+1_, conditioned on the path observed so far:

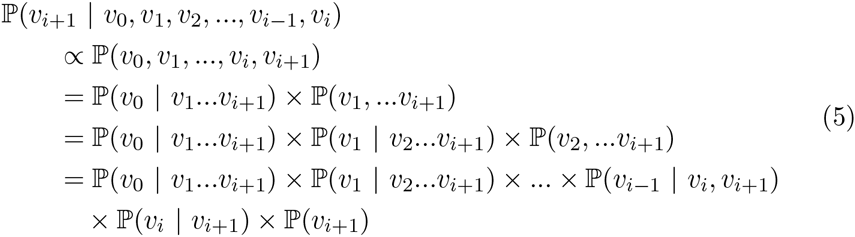

### 1.3 Simplification of conditional edge weights

Clearly, the number of factors in Equation 5 increases with *i*. For longer paths (more single nucleotide polymorphisms detected along the target region of interest), evaluating the equation becomes more computationally expensive, and risks potentially compounding estimation errors.

To construct a whole path *p* from *υ*_1_…*υ_m_*, the upper bound for the number of iterations will be |Σ| × *m* with calculations becoming increasingly complex as *i* increases.

To reduce complexity, we make an assumption of conditional independence between variants. Whilst this seems counter intuitive, the Naive Bayes model can deliver robust results despite its coarse assumption.

Thus we may simplify our previous equation and consider only the pairwise appearances of each *υ_i_* encountered thus far against *υ*_*i*+1_.

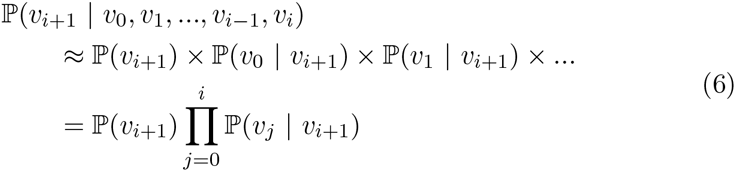

However as discussed in our article, reads will not cover all SNP positions 1.. *m* (if they did, we would not have to define this problem). Thus, we need not consider all variants in the current path when evaluating edge weights. Instead, we could limit the number of variants to consider, starting from the head of the path:

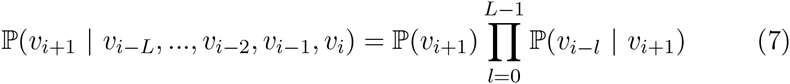

We define *L* as the the ‘lookback’ size, the number of variants of the current path to consider when selecting *υ*_*i*+1_. Conveniently, there is a reasonable intuition available for selecting a value for *L*: the mean number of SNP sites covered by the observed reads. Thus we avoid the scenario of introducing an algorithmically influential but mysterious parameter, such as *k*-mer size for metagenomic assembly.

### 1.4 Estimation of pairwise conditional probabilities

We must still devise a method to calculate the components of Equation 7. We present the following approximations, inspired by the Bag of Words model, commonly implemented in text classification domains (such as a spam filters).

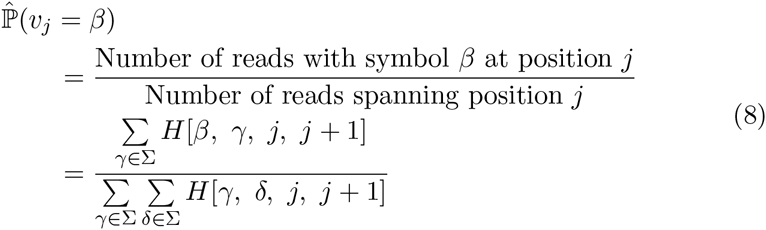

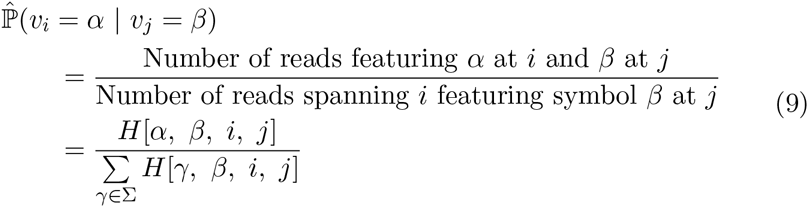

### 1.5 Smoothing

To avoid the possibility of dividing by 0 in cases of Equation 9 where a suitable read spanning *i* and *j* = *β* does not exist, we apply Laplace smoothing to effectively add a dummy support read. Future work will investigate alternative smoothing methodology.

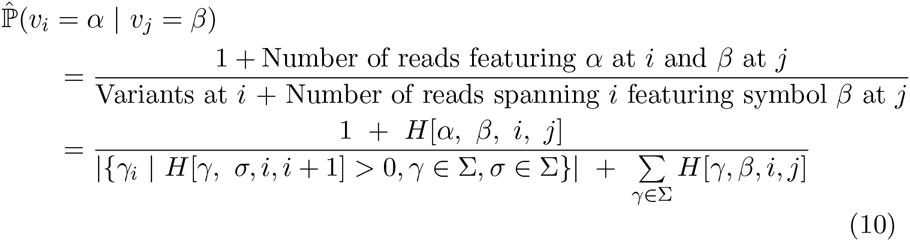

### 1.6 Re-weighting

The paths generated by `Gretel` are probabilistic, but not stochastic. For a given *H*, `Gretel` will always return the same path. Since it is the elements of *H* that effectively drive traversal of *G*, we can perform some form of post-path generation transformation of *H* to prevent repetitive generation of the same path and return the next most likely path on the next iteration instead.

Given a path *p*, we inspect the marginal distribution of each variant for all *i* ∈ 1..*m* (*i.e*. the probability of selecting the same variant if we were looking at the site in isolation), and determine the smallest marginal. `Gretel` iterates over each variant *p*[*i*] in the path, and uses the Hansel interface to re-weight the element *H*[*p*[*i*],*p*[*i* + 1],*i,i* + 1] by subtracting the result of multiplying the smallest marginal by the original value for the observation:

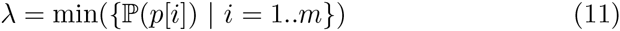

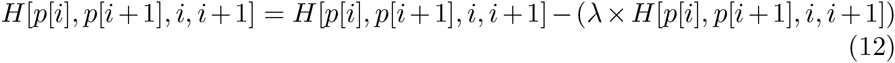

## Footnotes

1 A phrase that did not seem to catch on in the literature

